# Cdk9, Spt5 and histone H2B mono-ubiquitylation cooperate to ensure antisense suppression by the Clr6-CII/Rpd3S HDAC complex

**DOI:** 10.1101/240135

**Authors:** Miriam Sansó, Daniel Pinto, Peter Svensson, Viviane Pagé, Pabitra Parua, Danny A. Bitton, Jean Mbogning, Patricia Garcia, Elena Hidalgo, François Robert, Jürg Bähler, Jason C. Tanny, Robert P. Fisher

## Abstract

Cyclin-dependent kinase 9 (Cdk9) and histone H2B monoubiquitylation (H2Bub1) are both implicated in elongation by RNA polymerase II (RNAPII). In fission yeast, Cdk9 and H2Bub1 regulate each other through a feedback loop involving phosphorylation of the elongation factor Spt5. Conversely, genetic interactions suggest opposing functions of H2Bub1 and Cdk9 through an Spt5-independent pathway. To understand these interactions, we performed RNA-seq analysis after H2Bub1 loss, Cdk9 inhibition, or both. Either Cdk9 inhibition or H2Bub1 loss increased levels of antisense transcription initiating within coding regions of distinct subsets of genes; ablation of both pathways led to antisense derepression affecting over half the genome. Cdk9 and H2Bub1 cooperate to suppress antisense transcription by promoting function of the Clr6-CII histone deacetylase (HDAC) complex. H2Bub1 plays a second role, in opposition to Clr6-CII, to promote sense transcription in subtelomeric regions. Therefore, functional genomics revealed both collaborative and antagonistic functions of H2Bub1 and Cdk9.

## Introduction

Regulation of transcription elongation by RNA polymerase II (RNAPII) determines expression levels of many genes, including ones needed for cell-division and developmental-fate decisions in multicellular eukaryotes (1). Among the factors that directly influence transcription by promoting or impeding elongation in metazoans are key regulators of cell growth, proliferation and differentiation, including oncogene and tumor suppressor gene products. Factors involved in RNA processing and chromatin modification are recruited directly to the RNAPII elongation complex, ensuring that these events are kinetically coupled to RNA synthesis (2). Despite a growing inventory of proteins and protein modifications implicated in elongation control, many of the underlying molecular mechanisms are still incompletely understood.

Mono-ubiquitylation of histone H2B (H2Bub1) occurs in concert with elongation by RNAPII, and is catalyzed by the E2 ubiquitin conjugating enzyme Rad6 and E3 ubiquitin ligases related to *S. cerevisiae* Bre1 (3). Consequences of ablating the mammalian Bre1 orthologs RNF20 and RNF40 in cell lines or in vivo suggest important roles for H2Bub1 in regulating cell differentiation, migration and growth. RNF20, moreover, is overexpressed or silenced in different patient-derived tumor samples (4-6). H2Bub1 positively regulates the methylation of histone H3 at Lys4 (H3K4) and Lys79 (H3K79)—marks also associated with transcribed chromatin (7-9)—but H2Bub1 also functions independently of histone methylation (10,11). Methylation-independent functions of H2Bub1 include regulation of nucleosome stability and positioning within coding regions (12-14). The mechanisms for these effects and their consequences for gene expression are not known. Despite the presence of H2Bub1 at every transcribed gene, mutations that abolish H2Bub1 affect steady-state levels of only a small fraction of mRNAs, possibly suggesting redundant or compensatory homeostatic mechanisms (4,10,12,15).

In yeast and metazoans, co-transcriptional formation of H2Bub1 depends on Cdk9, an essential CDK that promotes elongation, helps recruit RNA-processing and histone-modifying enzymes to the transcription complex, and can directly influence the activity of those enzymes (16-18). In metazoans, Cdk9 is the catalytic subunit of positive transcription elongation factor b (P-TEFb), which works to overcome promoter-proximal pausing by RNAPII. Cdk9 executes its diverse functions in part through phosphorylation of the elongation factor Spt5. Spt5 phosphorylation creates a binding site for Rtf1, an important cofactor for H2Bub1 deposition (19-21). In the fission yeast *Schizosaccharomyces pombe*, Rtf1 and the Bre1 ortholog Brl2 promote Cdk9 association with chromatin and phosphorylation of Spt5 in a positive feedback loop (21,22). An analogous feedback mechanism involving Cdk9 and RNF20 operates in mammalian cells (23). In *S. pombe*, combining a mutation that ablates the major sites of Cdk9-dependent phosphorylation in Spt5 with deletion of *set1^+^*, which encodes the methyltransferase responsible for H3K4me, recapitulated the morphological and growth phenotypes associated with H2Bub1 loss in an H2Bub1-proficient strain (22). Cdk9 therefore acts both upstream and downstream of H2Bub1, through Spt5, to influence gene expression.

There is also genetic and genomic evidence in *S. pombe*, however, indicating functional opposition between Cdk9 and H2Bub1. For example, ablation of the H2Bub1 pathway or impairment of Cdk9 had opposite effects on RNAPII distribution within gene bodies: loss of H2Bub1 caused a shift in occupancy towards 3’-ends of genes, whereas a hypomorphic allele of *cdk9* caused RNAPII accumulation towards 5’-ends and depletion from downstream regions. Moreover, deletion of *brl2^+^* or removal of the histone H2B ubiquitin acceptor site by an *htb1-K119R* mutation produced cell morphology and septation defects that were reversed upon inactivation of Cdk9 (22). This suppression was not recapitulated by preventing Spt5 phosphorylation, indicating that antagonism between H2Bub1 and Cdk9 involves at least one Cdk9 target other than Spt5. Therefore, different Cdk9-dependent pathways are likely to underlie the reinforcing and opposing roles of Cdk9 and H2Bub1.

In order to define more clearly the relationships between Cdk9 and H2Bub1 in global gene expression, we performed strand-specific RNA-seq in *S. pombe* cells deficient in H2Bub1, Cdk9 activity or both. The most prevalent changes in RNA steady-state levels we detected, due to loss of either H2Bub1 or Cdk9 activity, were increases in antisense transcripts derived from protein-coding genes. When both pathways were impaired, antisense transcription increased at the majority of genes. Cdk9 and H2Bub1 acted synergistically to regulate antisense transcription by promoting the function of the Clr6-CII HDAC complex in gene coding regions. H2Bub1 also plays a Cdk9-independent role in activation of subtelomeric genes by opposing the silencing function of the same HDAC. These results reveal independent, collaborative or antagonistic effects at different genes transcribed by RNAPII and in distinct regions of the genome, which might account for the complex genetic interactions between Cdk9 and H2Bub1.

## Results

### Cdk9 and H2Bub1 collaborate to suppress antisense transcription

Previous microarray analyses in *S. pombe* demonstrated selective, largely non-overlapping effects of Cdk9 inhibition and H2Bub1 ablation on expression of protein-coding genes (10,24). To extend these analyses and uncover effects on global gene expression that might explain genetic interactions between Cdk9 and H2Bub1 pathways, we performed strand-specific RNA-seq analysis comparing patterns of transcript accumulation between wild-type cells and ones with 1) a *cdk9^as^* mutation that confers sensitivity to inhibition by the bulky adenine analog 3-MB-PP1 (24), 2) *htb1-K119R* or 3) both mutations. Cdk9 inhibition or H2Bub1 ablation had gene-specific rather than global effects on steady-state mRNA abundance (Fig. 1a), consistent with previous microarray analyses (10,24). We also detected 3-MB-PP1-dependent increases in antisense and intergenic transcript abundance in *cdk9^as^* cells, and drug-independent increases of similar magnitude in *htb1-K119R* cells. Finally, the *cdk9^as^ htb1-K119R* mutant resembled the *htb1-K119R* single mutant in the absence of drugs but, upon addition of 3-MB-PP1, had the highest levels of antisense and intergenic transcripts of any of the four strains. Individually, allele-specific inhibition of Cdk9 or a mutation blocking H2Bub1 affected sense transcript levels from <5% of *S. pombe* genes but led to increased antisense transcript levels from ~10% of genes (Fig. 1b and Supplementary Table 1). There was no significant overlap between the transcripts affected in each of the single-mutant strains (sense or antisense) with the exception of the increased antisense transcripts, for which the common set comprised ~25% of genes affected by either mutation alone (*P*<10^−15^; hypergeometric test)(Fig. 1c). Inactivation of Cdk9 in an *htb1-K119R* background led to increased sense transcript accumulation from ~5% of genes, whereas antisense transcripts were increased from >50% of genes (Fig. 1b). The latter group comprised all 135 genes similarly affected by either *htb1-K119R* or Cdk9 inhibition, most of those affected in one case but not the other, and nearly 2,000 additional genes that were not affected in either single-mutant strain (Fig. 1c). Quantification of individual transcripts revealed synergistic effects of Cdk9 inhibition and H2Bub1 loss at genes where antisense transcription was induced by either condition individually; median induction was ~4-fold over wild-type in *htb1-K119R* or 3-MB-PP1-treated *cdk9^as^* cells, but ~12-fold over wild-type in the drug-exposed double mutant (Fig. 1d). Therefore, whereas distinct subsets of genes have non-redundant requirements for Cdk9 or H2Bub1, the majority of protein-coding genes depend on wild-type function of either Cdk9 or the H2Bub1 pathway to suppress antisense transcription.

**Figure 1.**
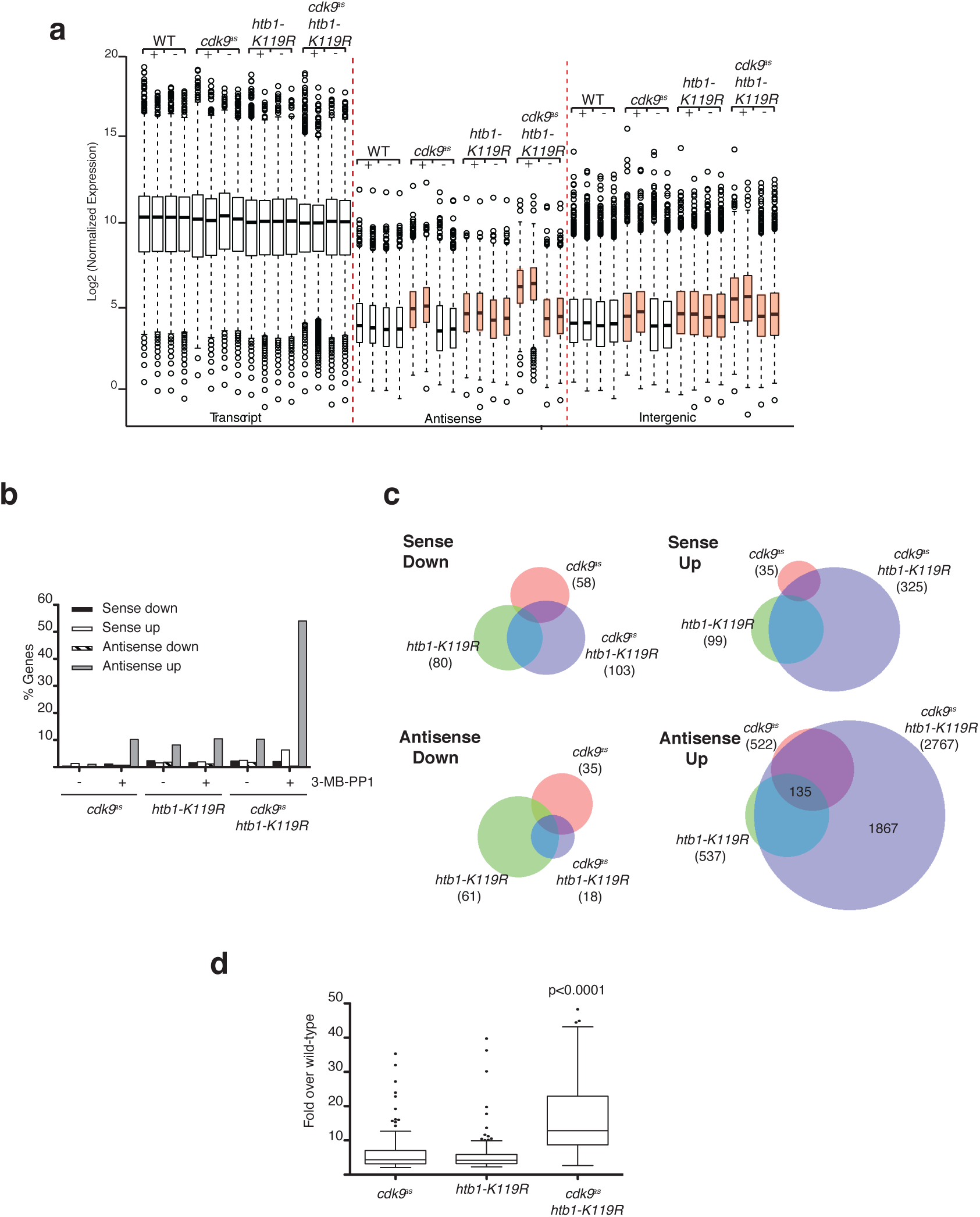
Cdk9 inhibition and *htb1-K119R* synergistically increase antisense transcripts throughout the genome. **(a)** Box-and-whisker plots of normalized expression levels for transcripts in the indicated categories for the indicated strains (JTB362, JTB299, JTB86, JTB508). “^+^” denotes treatment with 3-MB-PP1; “−” denotes treatment with DMSO. Orange boxes highlight increased levels of antisense and intergenic transcripts relative to wild-type. **(b)** Plot of the percentage of protein-coding genes in the indicated categories, defined by a minimum 2-fold change from the wild-type expression level in each condition (see Materials and Methods). **(c)** Venn diagrams comparing the differentially expressed transcripts (>2-fold change from wild-type) for the indicated strains in the presence of 3-MB-PP1. Numbers in brackets indicate total number of differentially expressed transcripts in the indicated strain. **(d)** Box-and-whisker plot of the fold-induction (in the presence of 3-MB-PP1) for 135 antisense transcripts regulated by Cdk9 activity and H2Bub1. Significance of the increased antisense expression levels in the double mutant was determined using the Mann-Whitney test.

We validated the RNA-seq results at *cdc2^+^*, a gene whose antisense transcript was induced by Cdk9 inhibition or *htb1-K119R* alone, and *erg32^+^*, one whose antisense transcript was only up-regulated by combined Cdk9 and H2Bub1 ablation. Strand-specific RT-qPCR confirmed modest effects of either Cdk9 inhibition or *htb1-K119R* on antisense transcript levels from *cdc2^+^* (relative to sense transcript levels at the control *act1^+^* locus), and a larger increase in antisense transcripts when both Cdk9 activity and H2Bub1 were blocked. The effect was more clearly synergistic at *erg32^+^*, where either single mutation alone had only minimal effects on antisense levels (Fig. 2a). For *cdc2^+^* we confirmed that the antisense transcript we detected corresponded to a previously annotated non-coding RNA, by RT-PCR with a primer pair specific for transcripts that begin within the gene body but extend upstream of the TSS (Supplementary Fig. 1a), and by northern blot hybridization with a strand-specific probe (Supplementary Fig. 1b).

**Figure 2.**
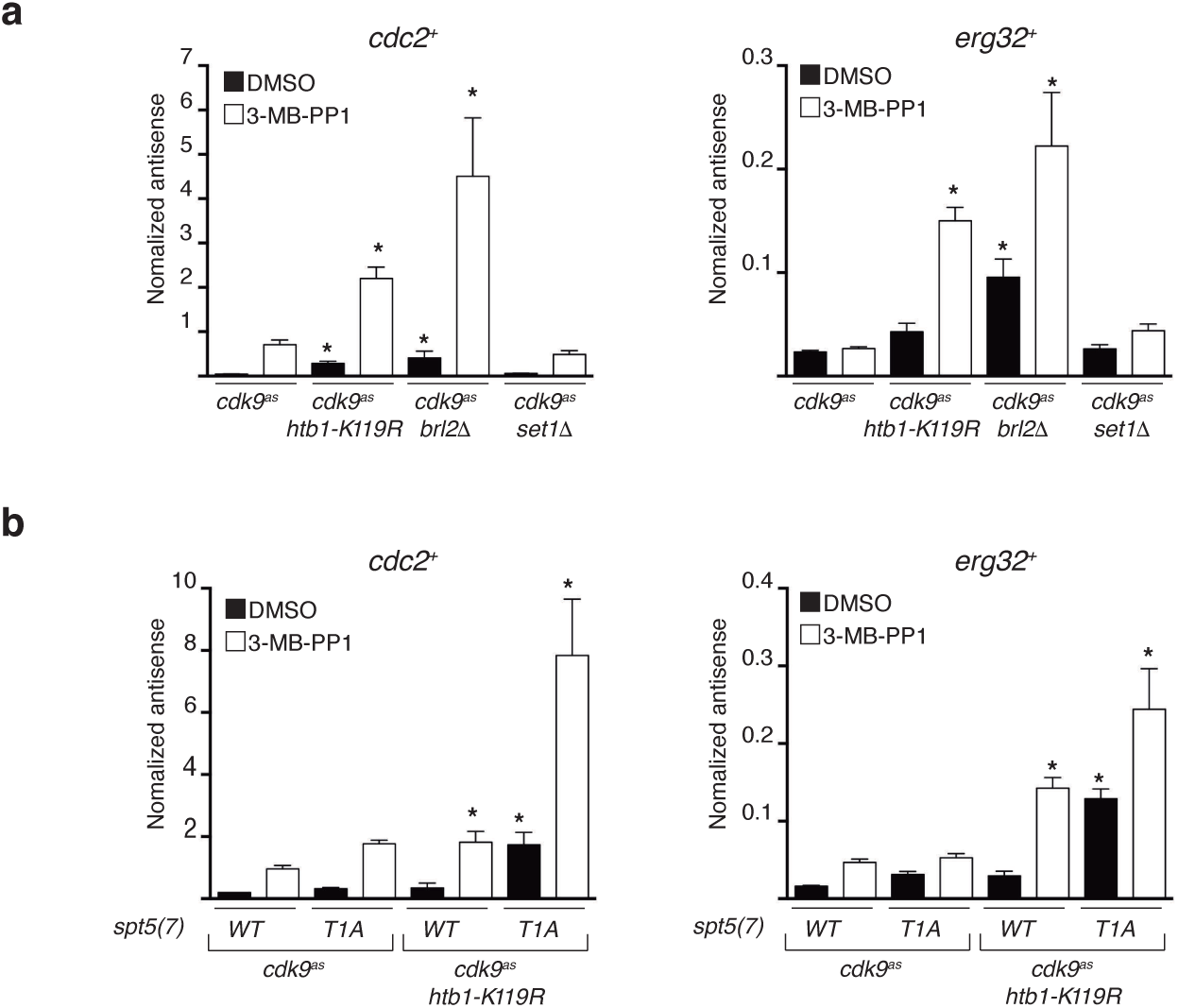
Antisense induction by Cdk9 inhibition and *htb1-K119R* is independent of histone H3 lysine 4 methylation and is enhanced by *spt5-T1A*. **(a)** Antisense transcript levels at the *cdc2^+^* and *erg32^+^* loci were quantified by strand-specific RT-qPCR in the indicated strains (JTB425, JTB508, JTB386, JTB408, respectively). Asterisks denote significant differences (*p*<0.05; unpaired t-test) from the DMSO-treated *cdk9as* strain (n=3; S.D.). (b) As in (a) for the indicated set of strains (JTB443, JTB444, JTB738–2, JTB736, respectively).

### Antisense suppression by Cdk9 and H2Bub1 is independent of H3K4 methylation and involves multiple Cdk9 targets

To determine if derepression of antisense transcription by the *htb1-K119R* mutation was due to loss of H2Bub1 per se, we measured levels of sense and antisense *cdc2^+^* and *erg32^+^* transcripts in a *cdk9^as^brl2Δ* strain by RT-qPCR. As was the case for *htb1-K119R*, a *brl2Δ* mutation enhanced antisense transcript levels induced by Cdk9 inhibition at both loci, indicating that antisense dysregulation by *htb1-K119R* is due specifically to loss of ubiquitylation, rather than another effect of the Lys➔Arg mutation (Fig. 2a). Indeed, compared to *htb1-K119R*, loss of the E3 led to higher levels of antisense transcription at both genes when Cdk9 was inhibited, and produced significantly more antisense derepression at *erg32^+^* in the absence of 3-MB-PP1. Therefore, Brl2 might have additional targets or functions that suppress antisense transcription. We also examined the role of the H3K4 methyltransferase Set1, activity of which is positively regulated by H2Bub1, in antisense regulation. The *set1Δ* mutation did not influence antisense transcript levels at either gene, arguing that antisense regulation by H2Bub1 is independent of downstream H3K4 methylation (Fig. 2a).

To understand how Cdk9 suppresses antisense transcription, we sought to identify its relevant target(s). Spt5 is a major substrate of *S. pombe* Cdk9: the majority of phosphorylation occurs within a carboxy-terminal domain (CTD) comprising 18 repeats of a nonapeptide motif (consensus: T_1_P_2_A3W4N_5_S_6_G7S_8_K_9_). Within the repeats, Cdk9 phosphorylates the Thr1 position exclusively (25), and inhibition of Cdk9 in vivo abolishes detectable phosphorylation of this site (22). We analyzed antisense transcript levels in strains expressing Spt5 variants with only seven repeats—a truncation that did not by itself compromise cell growth or viability (26). Although a mutation that removed Thr1 from each repeat—*spt5*(*T1A*)*_7_–did* not increase antisense transcription of *cdc2^+^* or *erg32^+^* in an *htb1^+^* strain with active Cdk9, it produced levels of antisense transcripts similar to those caused by Cdk9 inhibition in an *htb1-K119R* background (Fig. 2b). Moreover, *spt5*(*T1A*)*_7_* strains remained sensitive to Cdk9 inhibition; treatment of *cdk9^as^ htb1-K119R spt5*(*T1A*)_7_ cells with 3-MB-PP1 produced the highest levels of antisense transcripts detected at either gene. Therefore, constitutive loss of Spt5 Thr1 phosphorylation (Spt5-T1P) is not sufficient to derepress antisense transcription of *cdc2^+^* or *erg32^+^* in H2Bub1-proficient cells, but sensitizes cells to antisense induction by loss of H2Bub1, *and* to combined H2Bub1 loss and Cdk9 inhibition (which presumably involves at least one other Cdk9 target).

### Cdk9 and H2Bub1 promote HDAC function to suppress antisense transcription

Previous studies in fission yeast identified two classes of factors involved in regulation of antisense transcription: ones that work by regulating chromatin structure, such as the Clr6-CII HDAC complex, orthologous to the budding yeast Rpd3S complex that suppresses cryptic transcription initiation (sense and antisense) within genes transcribed by RNAPII (27); and those that target antisense transcripts for post-transcriptional elimination by the nuclear exosome (28-31). To ask if either of these mechanisms was involved in antisense suppression by Cdk9 and H2Bub1, we performed hierarchical clustering analysis to compare our antisense expression data with published profiles obtained in various *S. pombe* mutants (Fig. 3a). These analyses revealed four clusters of highly correlated (ρ≥0.5) expression patterns. As expected, mutants affecting co-transcriptional (“chromatin cluster”) or post-transcriptional (“exosome cluster”) antisense regulation formed distinct groups. Data sets derived from strains harboring a *cdk9^as^* allele clustered together in a group distinct from those from strains with *htb1-K119R* alone. The *cdk9^as^* cluster includes data sets from control (DMSO-treated) samples, likely reflecting an influence of the mildly hypomorphic *cdk9^as^* allele on antisense regulation even in the absence of the inhibitory analog. The 3-MB-PP1-treated *cdk9^as^ htb1-K119R* samples were more strongly correlated with the chromatin cluster than were the DMSO-treated samples (Fig. 3a; compare dark to light blue in the highlighted portion of the heatmap), suggesting a close relationship between the patterns of antisense transcription due to combined Cdk9 inhibition and H2Bub1 loss and those caused by defects in chromatin structure.

**Figure 3.**
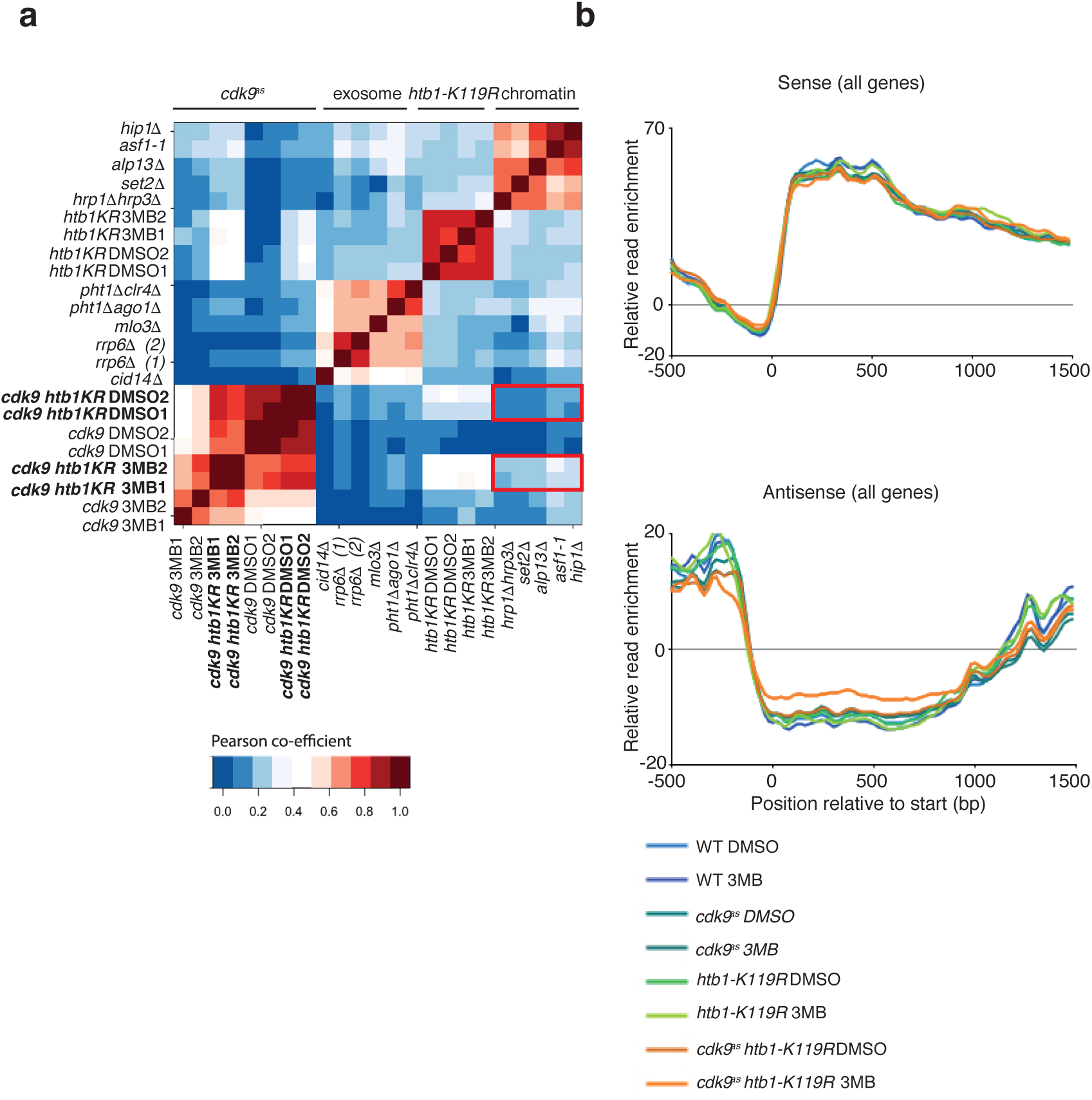
Antisense induction by Cdk9 inhibition and *htb1-K119R* is related to that caused by defects in chromatin-modifiying factors. **(a)** Heat map showing correlations between global patterns of antisense induction in the indicated datasets. Red boxes highlight differences in correlation between the *“cdk9^as^”* cluster and the “chromatin” cluster. **(b)** Metagene profiles of sense and antisense transcripts averaged across 5,123 protein-coding genes in the indicated strains. X-axis denotes position relative to transcription start site on an average gene.

Metagene analysis of the antisense transcripts induced in the *cdk9^as^ htb1-K119R* strain revealed that the signals were uniformly distributed throughout gene coding regions (Fig. 3b). Therefore, most of these transcripts arise from within the bodies of transcribed genes, consistent with a functional link to Clr6-CII and related factors (32-34). There was no apparent bias, in any of the conditions tested, towards adjacent gene pairs that are transcribed convergently, which might give rise to antisense transcripts due to inefficient termination leading to read-through transcription, further supporting an intragenic origin of these transcripts (Supplementary Fig. 2a-c).

### Genetic interactions between *cdk9* or *htb1* and the Clr6-CII complex

Consistent with a correlation between changes in *cdk9^as^ htb1-K119R* cells and other mutants defective in chromatin regulation, *cdc2^+^* and *erg32^+^* antisense transcript levels were increased by deletion of *cph1^+^* or *alp13^+^*, which encode subunits of Clr6-CII, or of *set2^+^*, which encodes a histone H3 Lys36 (H3K36) methyltransferase that promotes Clr6-CII function (Fig. 4a and Supplementary Fig. 3a)(28,29). Deletion of *hrp3^+^*, encoding a chromatin remodeling factor thought to regulate antisense transcription, also increased antisense transcription levels at these loci (32,35,36). Antisense transcription at either gene was unaffected by *hos2Δ*, which ablates Set3C, another HDAC implicated in antisense suppression (37), suggesting that the increases we detected are likely to reflect impaired function of a specific HDAC rather than a general decrease in deacetylase activity. Therefore, Set2, Clr6-CII, and Hrp3 are candidate effectors of the antisense-suppression functions of Cdk9 or H2Bub1.

**Figure 4.**
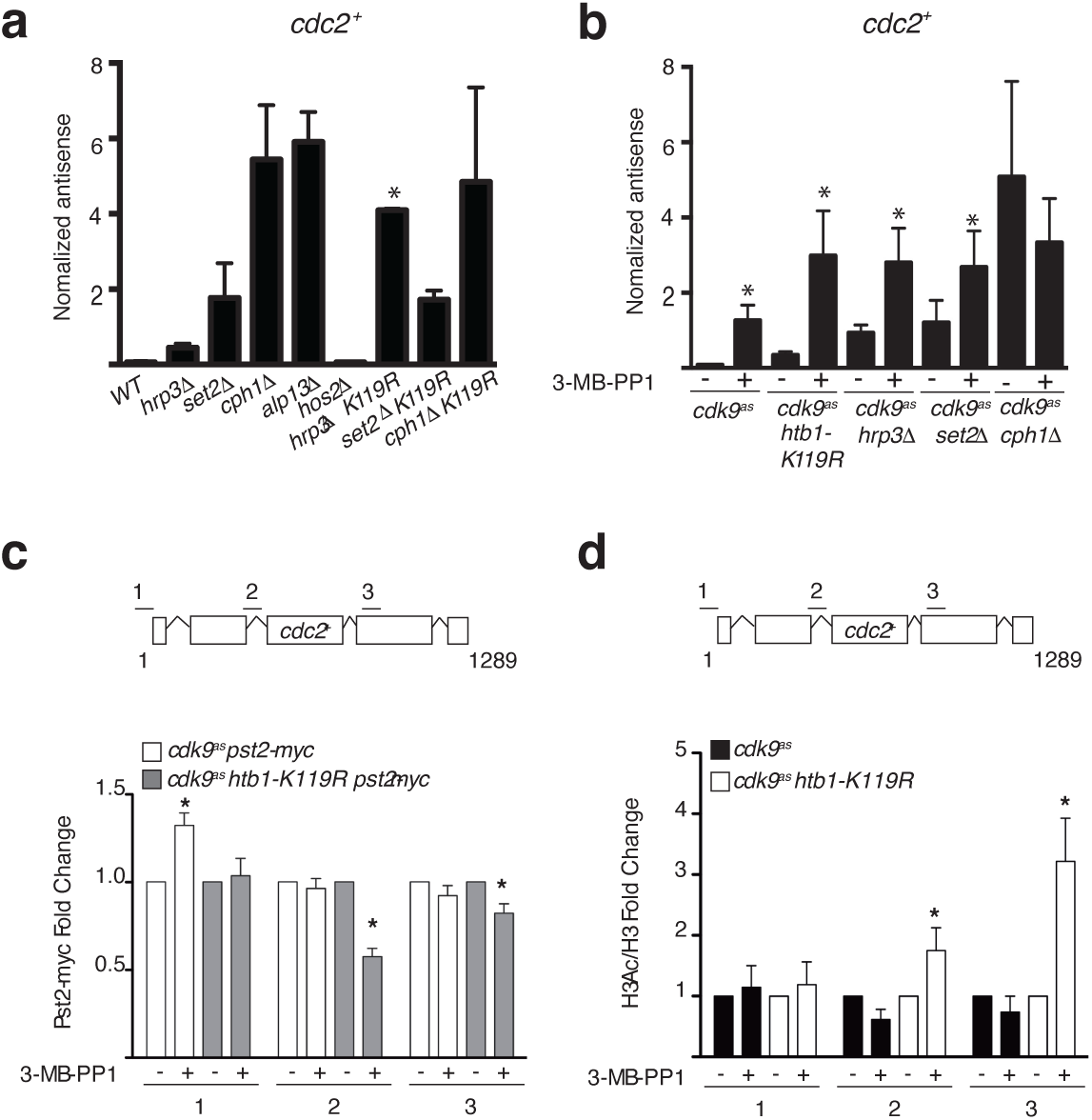
Cdk9 and H2Bub1 regulate the Clr6-CII HDAC complex. **(a)** Antisense transcript levels at *cdc2^+^* in the indicated strains (JTB362, JTB636, JTB142, JTB414, JTB413, JTB277, JTB618, JTB154, JTB640, respectively) measured by RT-qPCR (n=3; S.D.). Asterisks denote a significant difference from the single mutant strain (*p*<0.05; unpaired t-test). **(b)** Antisense transcript levels at *cdc2^+^* in the indicated strains (JTB425, JTB508, JTB639, JTB442, JTB505, respectively) in the presence or absence of 3-MB-PP1 measured by RT-qPCR (n=4; S.D.). Asterisks denote a significant difference between the presence and absence of 3-MB-PP1 (*p*<0.05; unpaired t-test). **(c)** ChIP of Pst2-myc was performed in the indicated strains (JTB838 and JTB842) treated with DMSO (−) or 20 μM 3-MB-PP1 (+). Signals were quantified by qPCR and normalized to the DMSO control for each strain. Diagram of the *cdc2^+^* locus indicating the positions of primer pairs used for ChIP-qPCR is indicated on top. * denotes a statistically significant difference from control (*p*<0.05; unpaired t-tests). **(d)** ChIP of acetylated H3 was performed in the indicated strains (JTB425 and JTB508) and normalized to total H3 occupancy. The H3Ac/H3 ratio was then normalized to the DMSO control for each strain. * denotes a statistically significant difference from control (*p*<0.05; unpaired t-tests).

To analyze these relationships in more detail, we measured *cdc2*^+^ and *erg32*^+^ antisense levels in a series of double mutant strains. At both genes, an *htb1-K119R* mutation enhanced antisense transcription in combination with *hrp3Δ*, arguing that H2Bub1 and Hrp3 function in parallel pathways. In contrast, antisense induction by *htb1-K119R* was not additive with that of *set2Δ* or *cph1Δ*, suggesting genetic epistasis (Fig. 4a and Supplementary Fig. 3a). Inhibition of Cdk9 induced antisense transcription at *cdc2^+^* in an *hrp3Δ* or *set2Δ*, but not a *cph1Δ*, background (Fig. 4b). We did not detect an additional effect of Cdk9 inhibition on *erg32^+^* antisense transcription in any of these mutants (Supplementary Fig. 3b). Taken together, the epistasis relationships suggest that Clr6-CII might be a common effector of antisense suppression by Cdk9 and H2Bub1.

### Cdk9 and H2Bub1 promote Clr6-CII chromatin association and activity

These results seemed to place H2Bub1 and Cdk9 upstream of a Clr6-CII function needed to suppress antisense transcription. To test for an effect of H2Bub1 loss or Cdk9 inactivation on Clr6-CII recruitment, we performed chromatin immunoprecipitation and quantitative PCR (ChIP-qPCR) analysis of the Clr6-CII subunit Pst2 in *pst2-myc* strains. At *cdc2^+^* and *erg32^+^*, there was no significant loss of Pst2-myc signal due to Cdk9 inhibition alone. However, Pst2-myc cross-linking was significantly decreased in the coding regions of both genes upon 3-MB-PP1 treatment of *cdk9^as^ htb1-K119R* cells (Fig. 4c and Supplementary Fig. 3c). No change in Pst2-myc ChIP signals was detected at *act1^+^*, where there was no increase in antisense transcripts (Supplementary Fig. 3d). Therefore, either Cdk9 activity or H2Bub1 is required for normal levels of Clr6-CII recruitment to coding regions of sensitive genes.

To test for effects of H2Bub1 or Cdk9 on Clr6-CII *function*, we measured histone H3 acetylation in *cdk9^as^* or *cdk9^as^ htb1-K119R* strains. Consistent with decreased Clr6-CII occupancy, histone H3 acetylation at Lys9 and Lys14, normalized to total H3 levels, was significantly increased within coding regions of both *cdc2^+^* and *erg32^+^* (Fig. 4d and Supplementary Fig. 3e), but not of *act1^+^* (Supplementary Fig. 3f), only when both H2Bub1 and Cdk9 activity were blocked. ChIP analysis in the Clr6-CII mutant strains *alp13Δ* and *cph1Δ*, and in the *set2Δ* strain, also revealed increased levels of histone acetylation at both loci relative to wild-type cells. In contrast, the effects of *hrp3Δ*. on acetylation were gene-specific, with increases at *erg32^+^* but not *cdc2^+^* (Supplementary Fig. 4a, b). Therefore, the increased levels of antisense transcripts detected in 3-MB-PP1-treated *cdk9^as^ htb1-K119R* cells correlated with impaired recruitment and function of the HDAC Clr6-CII.

### A specialized role of H2Bub1 in subtelomeric gene expression

In contrast to the widespread effects of combined Cdk9 inhibition and H2Bub1 loss on antisense transcription, ablation of either pathway alone led to more restricted, largely non-overlapping effects—repression and induction—of sense transcripts (Figs 1b, c and 2b). Given their known interactions with other factors that influence chromatin structure, we considered the possibility that either Cdk9 or H2Bub1 might affect transcription preferentially within specific sub-chromosomal domains or regions. We analyzed the RNA-seq data to discern any bias in the positions where changes occurred within the *S. pombe* genome. As expected, given the number of genes affected, increased antisense transcription occurred throughout the genome in all of the mutant strains. However, there was significant enrichment of telomere-proximal genes in the set of sense transcripts that decreased in *htb1-K119R* relative to wild-type cells (Fig. 5a, b). This bias did not appear in data from the *cdk9^as^* mutant—genes repressed by 3-MB-PP1 were relatively evenly distributed throughout the genome—but was largely maintained in a 3-MB-PP1-treated *cdk9^as^ htb1-K119R* double mutant (Fig. 5b).

**Figure 5.**
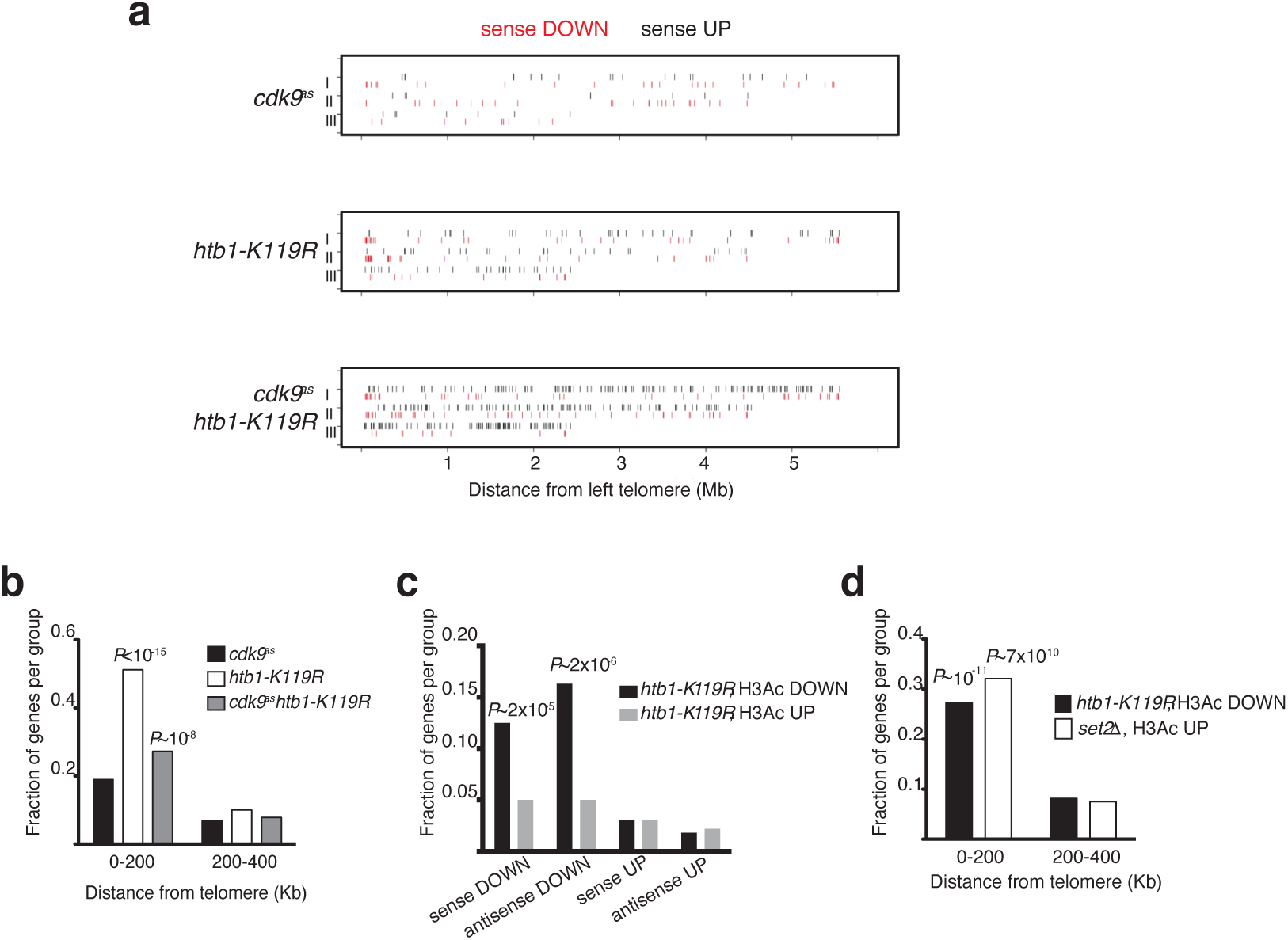
H2Bub1 is necessary for expression of subtelomeric genes. **(a)** Chromosomal distributions of differentially expressed sense transcripts (>2-fold change from wild-type) for the indicated strains in the presence of 3-MB-PP1. **(b)** Fraction of repressed genes for each strain within the indicated chromosomal intervals. Significance of enrichment was determined using the hypergeometric test. **(c)** Changes in histone acetylation at H2Bub1-regulated genes. Significance of enrichment was determined by the hypergeometric test. **(d)** Fraction of loci with the indicated H3 acetylation changes (1.5-fold or more compared to WT) within the indicated chromosomal intervals. Significance of enrichment was determined using the hypergeometric test.

Increased antisense transcription due to H2Bub1 loss and Cdk9 inhibition was associated with increased histone acetylation at affected loci (Fig. 4d and Supplementary Fig. 3e). To ask if the positional bias in genes positively regulated by H2Bub1 was similarly correlated with effects on histone acetylation, we performed ChIP followed by microarray hybridization analysis (ChIP-chip) to quantify levels of acetylated histone H3 (relative to total H3) at protein-coding and annotated non-coding RNA genes in wild-type and *htb1-K119R* strains. There was significant enrichment of hypoacetylated loci (defined by a ≥1.5-fold decrease relative to wild-type) among H2Bub1-dependent sense and antisense transcripts identified by RNA-seq (Fig. 5c). In contrast, there was no significant overlap between H2Bub1-repressed transcripts (sense or antisense) and loci hyperacetylated in *htb1-K119R*, perhaps consistent with the need to impair both Cdk9 and H2Bub1 in order to affect HDAC function. Hypoacetylated loci were also enriched within 200 kb of telomeres in *htb1-K119R* cells, supporting a Cdk9-independent role for H2Bub1 in promoting histone acetylation and transcription at subtelomeric genes (Fig. 5d).

Genes proximal to telomeres are subject to heterochromatic gene silencing, so a possible explanation for their decreased expression in *htb1-K119R* cells is inappropriate spreading of heterochromatin. However, genome-wide analyses did not detect increased H3K9 methylation—a marker of heterochromatin—proximal to telomeres in the *htb1-K119R* mutant (38). Moreover, deletion of *clr3^+^* or *sir2^+^*, which encode HDACs essential for heterochromatic silencing (39,40), did not cause elevated expression of the subtelomeric *aes1^+^* gene, and a *sir2Δ* mutation did not alleviate *aes1^+^* repression by *htb1-K119R* (Supplementary Fig. 5a).

A recent study showed that subtelomeric chromatin on chromosomes I and II is maintained in a highly condensed state by Set2 and Clr6 (41). By ChIP-chip analysis, histone H3 at subtelomeric loci was significantly hyperacetylated in a *set2Δ* strain (Fig. 5d), with significant overlap between subtelomeric genes hypoacetylated in the *htb1-K119R* strain and those hyperacetylated in *set2Δ* cells (*P*~9×10^−4^; hypergeometric test). This overlap was particularly striking within an ~60 kb window near *tel2L*, in which 14 of 27 genes showed opposite effects of *htb1-K119R* and *set2Δ* on acetylation (Fig. 6a).

**Figure 6.**
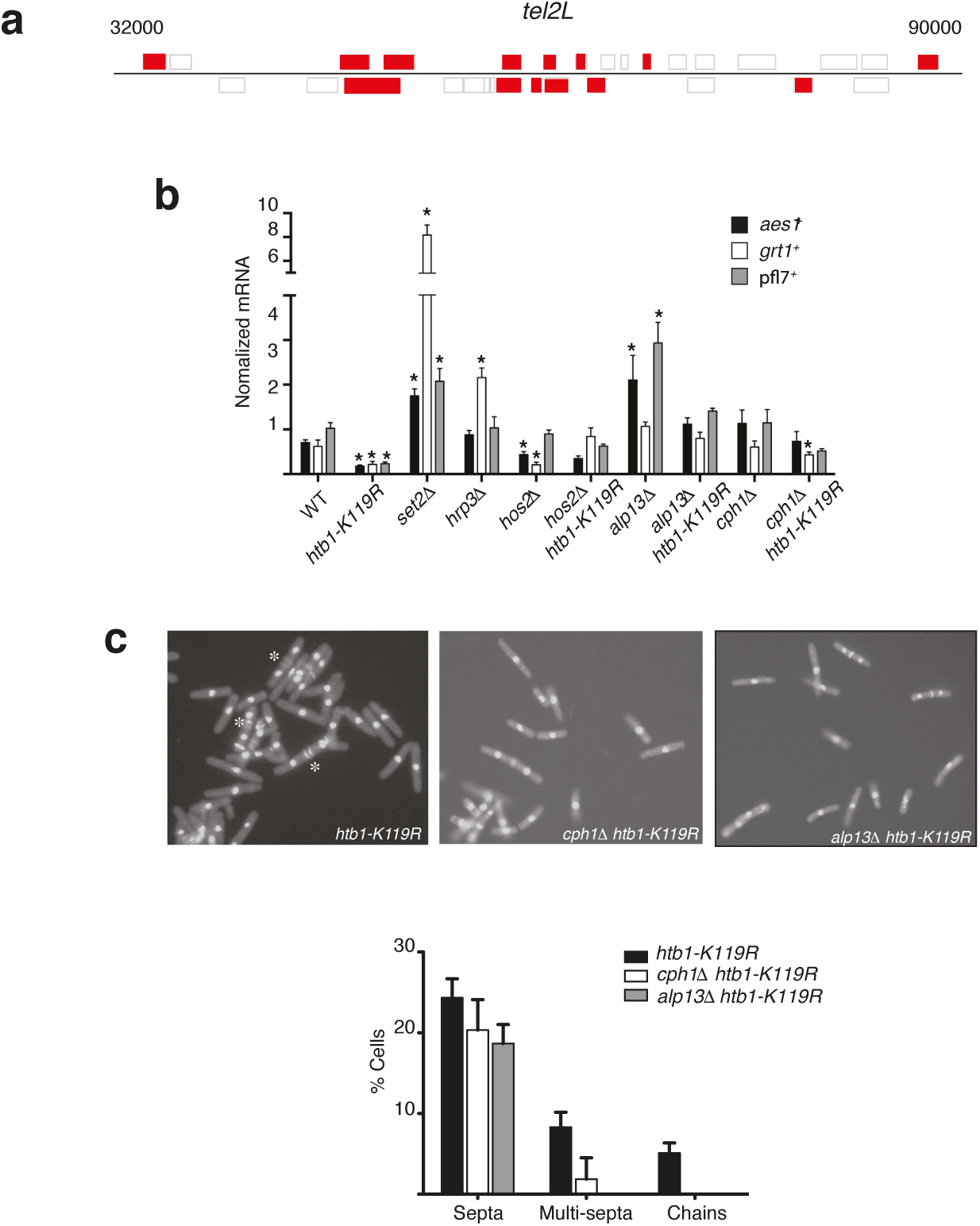
Evidence for opposing functions of H2Bub1 and the Clr6-CII HDAC complex. (a) Map of genes within a ~60 kb interval proximal to the left telomere of chromosome 2 (*tel2L* locus). Genes highlighted in red are those at which opposite effects on H3 acetylation were detected by ChIP in *htb1-K119R* and *set2Δ* strains. **(b)** Normalized mRNA expression levels of *aes1^+^, grt1^+^* and *pf17^+^* (n=3–5; S.D.), for the indicated strains (JTB362, JTB97, JTB142, JTB413, JTB637, JTB414, JTB640, respectively). **(c)** Top: fluorescent images of DAPI/calcofluor-stained cells from indicated strains (JTB97, JTB640, JTB637, respectively). Bottom: quantification of abnormal septation in strains from top panel (at least 200 cells counted per experiment, n=2).

To probe the antagonistic effects of H2Bub1 and Set2 at subtelomeric genes further, we measured transcript levels by RT-qPCR for *aes1^+^, grt1^+^*, and *pfl7^+^* genes, all located proximal to *tel2L*, in a panel of mutant strains. In agreement with our genome-wide analyses, all three genes were strongly repressed by *htb1-K119R* (Fig. 6b). Analysis of *brl2Δ* and *set1Δ* mutants confirmed that repression was due to loss of H2Bub1 per se, and not recapitulated by downstream loss of H3K4me (Supplementary Fig. 5b). In contrast, all three genes were induced by *set2Δ*, and two of three were significantly induced by *alp13Δ*, consistent with opposing functions of H2Bub1 and Clr6-CII at these genes (Fig. 5f). Furthermore, decreases in subtelomeric transcripts due to *htb1-K119R* were reversed in *alp13Δ htb1-K119R* and *cph1Δ htb1-K119R* double mutants. Therefore, whereas H2Bub1 works with Cdk9 to *promote* Clr6-CII function and suppress antisense transcription in gene bodies, H2Bub1 also acts independently of Cdk9 to *oppose* this HDAC and ensure normal levels of sense transcription in subtelomeric regions.

### Opposing effects of Cdk9 inhibition and H2Bub1 loss detected by RNA-seq

Our analysis thus far revealed complex interactions between H2Bub1 and Cdk9, including instances in which the two pathways acted either synergistically or antagonistically. Because we previously detected mutual genetic suppression suggestive of functional antagonism between Cdk9 and H2Bub1, we queried the RNA-seq data for sense or antisense transcripts that were affected by loss of H2Bub1 and either unaffected, or affected in the opposite direction, by 3-MB-PP1 treatment of a *cdk9^as^ htb1-K119R* strain. There were large fractions of sense (43%) and antisense (64%) transcripts repressed in the *htb1-K119R* mutant that met these criteria (Fig. 1c and Supplementary Table 2); subtelomeric genes were included but not significantly overrepresented among these genes (Supplementary Table 2). In contrast, only 11% of sense transcripts and 4% of antisense transcripts that were induced in the *htb1-K119R* mutant failed to be induced upon 3-MB-PP1 treatment of *cdk9^as^ htb1-K119R* (Fig. 1c). This suggests that opposing phenotypic consequences are more likely to stem from gene repression due to H2Bub1 loss that is rescued by Cdk9 inhibition. We reasoned that Cdk9 inactivation might impair Clr6-CII function, preventing repression and thereby alleviating phenotypic effects of *htb1-K119R*. Indeed, the cell morphology defects caused by *htb1-K119R*—increased frequencies of multi-septated and chained cells— were suppressed by mutations affecting Clr6-CII (Fig. 6c). This resembles the rescue of these phenotypes by Cdk9 inhibition (22). Taken together, our results indicate that Cdk9 and H2Bub1 pathways regulate the HDAC Clr6-CII in a context-dependent manner, influenced by genomic location (Fig. 7) and probably other factors. Moreover, the fact that genes can be differentially “wired” to respond positively or negatively to changes in H2Bub1 and/or Cdk9 activity might help to explain the complex genetic interactions between the two pathways.

**Figure 7.**
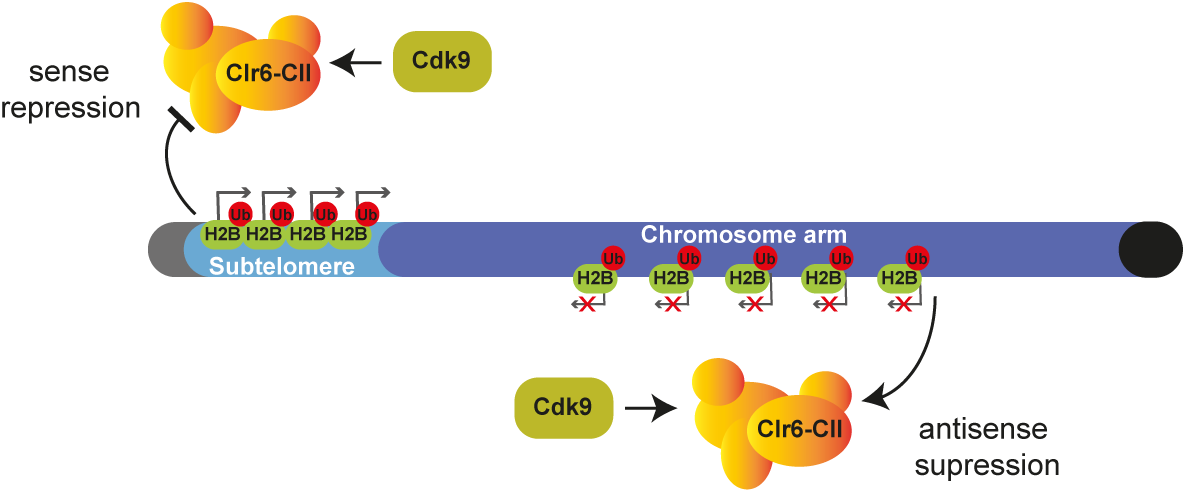
Model depicting interactions between Cdk9, H2Bub1, and the Clr6-CII HDAC on chromosome arms and at subtelomeric regions (see text for details).

## Discussion

Perturbations of chromatin structure due to passage of the transcription elongation complex (TEC) carry the inherent risk of inappropriate access by initiation factors. Accordingly, enzymes and factors that directly regulate elongation also work to suppress cryptic initiation of both sense and antisense transcripts through their influence on chromatin structure in gene coding regions (29). Here we have shown that antisense suppression at most RNAPII-transcribed genes in *S. pombe* requires either Cdk9 activity or H2Bub1 to promote the function of the Clr6-CII HDAC complex in coding regions.

Studies of the budding yeast Rpd3S complex—orthologous to *S. pombe* Clr6-CII—uncovered multiple determinants of its localization and function in gene coding regions. In vitro, Rpd3S preferentially binds to and deacetylates nucleosomes that harbor H3K36me, consistent with a functional link detected in vivo between this HDAC and the H3K36 methyltransferase Set2 (42). However, Rpd3S can interact with nucleosomes in the absence of histone modifications and is also influenced by linker DNA (42,43). Rpd3S association with transcribed genes can also occur independent of H3K36me in vivo, although H3K36me promotes Rpd3S-dependent histone deacetylation (44,45). Rpd3S binds to the phosphorylated Rpb1 CTD in vitro, and the *S. cerevisiae* orthologs of Cdk7 (Kin28) and Cdk9 (Bur1), as well as the Spt5 carboxy-terminal region, are implicated in Rpd3S recruitment to gene bodies (44,45). These data are in agreement with the potentiation of Rpd3S function by Bur1 and H3K36me detected by ChIP analysis at select genes (46). We have uncovered an analogous, cooperative effect involving Cdk9 and H2Bub1. H2Bub1 has not previously been linked directly to function of an Rpd3S-related HDAC, although the conserved MRG15 subunit (ortholog of Alp13) has been reported to interact with H2Bub1 in the context of the NuA4 histone acetyltransferase complex during DNA damage repair in mammalian cells (47). Our results suggest that H2Bub1 is an important feature of the chromatin landscape recognized by Rpd3S/Clr6-CII-type HDAC complexes. An emergent question is whether the Sin3B HDAC complex, the mammalian ortholog of Rpd3S and a potential therapeutic target in pancreatic cancer, is similarly responsive to Cdk9 or H2Bub1 (48,49).

Our data suggest that multiple Cdk9-dependent phosphorylation events contribute to antisense suppression, including Spt5-T1P. Spt5 was recently shown to prevent antisense transcription initiating within *S. pombe* genes, a function apparently linked to its role as an RNAPII processivity factor (50). Our results now implicate the Spt5 kinase Cdk9, and chromatin-modifying enzymes that work in concert with or downstream of Spt5—the H2Bub1 machinery and Clr6-CII—in global suppression of antisense transcription. Increased antisense transcription might arise through more than one Spt5-dependent mechanism; in the previous study the requirement in antisense suppression was uncovered by induced degradation of Spt5 protein, but not by a constitutive truncation removing the entire Spt5 CTD. In contrast, we show that specifically blocking CTD phosphorylation by *spt5*(*T1A*)*_7_* mutation induces antisense transcription in the absence of H2Bub1. To reconcile these results, we propose that the Spt5 CTD, when unphosphorylated, impedes elongation and favors antisense transcription—both functions lost when the CTD is removed—whereas the phosphorylated form promotes elongation and enhances H2Bub1 and HDAC function, thereby suppressing cryptic initiation within gene bodies.

The similar levels of antisense transcripts detected in H2Bub1-deficient cells when Cdk9 was inhibited or Spt5-Thr1 residues were changed to Ala suggest that Spt5-T1P is critical for the synergistic effect of Cdk9 inhibition and H2Bub1 loss. The lack of an effect due to *spt5*(*T1A*)*_7_* alone, however, argues that other Cdk9 targets can compensate for its absence in H2Bub1-proficient cells. Interestingly, the increased antisense transcription detected upon Spt5 depletion was largely restricted to the region in which RNAPII accumulated, within ~500 bp downstream of the sense TSS, suggesting a “checkpoint” after which Spt5 function is no longer needed either in productive elongation or in antisense suppression (50). In contrast, antisense transcripts arising due to combined H2Bub1 loss and Cdk9 inhibition were distributed throughout gene bodies, possibly reflecting loss of phosphorylation of other Cdk9 targets that act later in the transcription cycle.

Surprisingly, the *spt5*(*T1A*)*_7_* mutation *sensitized* H2Bub1-deficient cells to effects of Cdk9 inhibition on antisense transcription. Cdk9 inhibition renders Spt5-T1P undetectable in extracts (22), so an additive effect of *spt5*(*T1A*)*_7_* and Cdk9 inhibition might suggest a role of the wild-type CTD independent of its phosphorylation. Similarly, inhibition of Cdk9 makes H2Bub1 undetectable, but enhances antisense transcription in combination with an *htb1-K119R* mutation, possibly suggesting a function for Lys119 that is independent of H2Bub1. Alternatively, it is possible that Cdk9-independent pathways support some H2Bub1 in vivo, as suggested by the roles of H2Bub1 in DNA replication and repair—processes not likely to depend on Cdk9 activity (51,52).

In budding yeast, H2Bub1 enhances function of the HDAC Set3C, which is also implicated in antisense suppression (37,53). The effects we describe here, however, are independent of the *S. pombe* Set3C catalytic subunit Hos2. H2Bub1 has also been functionally linked to CHD-type ATP-dependent chromatin remodeling complexes (54). The *S. pombe* CHD-type remodelers repress antisense transcription by enforcing nucleosome positioning within gene bodies, a function also ascribed to H2Bub1 (12,36). At the loci we tested, antisense transcription increased upon deletion of *hrp3^+^*, the remodeler of this class that is most important for antisense regulation in *S. pombe*, but the effects were additive with either Cdk9 inhibition or H2Bub1 loss. This implies a function of Hrp3 parallel to, rather than downstream of, Cdk9 and H2Bub1.

Inhibition of Cdk9 and genetic ablation of H2Bub1 cause distinct effects on steady-state RNA levels, as seen in previous microarray analyses (10,24) and indicated here by 1) the largely non-overlapping sets of transcripts affected in *cdk9^as^* or *htb1-K119R* mutants, and 2) a Cdk9-independent role of H2Bub1 at subtelomeric genes. Decreased expression of those genes in *htb1-K119R* cells coincided with decreased levels of histone acetylation, the opposite of what occurred in *set2Δ* and Clr6-CII-deficient mutants. Set2 and Clr6 are involved in the formation of highly condensed “knob” structures comprising the subtelomeres of chromosomes I and II, which are spatially separate from telomeric heterochromatin (41). (The telomeres of chromosome III are adjacent to ribosomal DNA rather than canonical telomeric repeats, likely accounting for their distinct behavior.) Formation of these structures correlates with repression by Set2 and Clr6 (41,55), and we now show that these regions are associated with Set2-dependent histone hypoacetylation. Repression by *htb1-K119R* was alleviated by *alp13Δ* or *cph1Δ*, consistent with H2Bub1 and Clr6-CII having opposing functions at subtelomeric loci. Subtelomeric loci are also overrepresented among genes that depend on histone acetyltransferases (56) and chromatin remodeling complexes for expression (57). Subtelomeric chromatin shares features with the inner centromere, including the presence of dimethylated H3K4 and depletion of the histone H2A variant H2A.Z (58). H2Bub1 promotes transcription of non-coding RNAs from the inner centromere (38), suggesting that it could be part of a regulatory mechanism shared with subtelomeric genes.

Cdk9 promotes elongation of sense transcripts at most genes, and H2Bub1 facilitates both Cdk9 recruitment and Spt5-T1P (22-24,59,60). Here we show that Cdk9 and H2Bub1 work cooperatively, through the HDAC Clr6-CII, to repress intragenic transcription initiation in the majority of fission yeast genes—a function in which Spt5 is also implicated (50). At subtelomeric genes, however, H2Bub1 appears to play a positive role in transcription independent of Cdk9 and in opposition to Clr6-CII. Reducing activity of Cdk9 normalized expression of nearly half of the transcripts repressed by *htb1-K119R*, both proximal to telomeres and elsewhere in the genome. We propose that at these genes the predominant function of Clr6-CII, facilitated by Cdk9, is to repress transcription in opposition to H2Bub1; this antagonism might explain how inhibition of Cdk9 or loss of Clr6-CII subunits suppressed cell morphology phenotypes caused by H2Bub1 deficiency [(22); this report]. Consistent with this interpretation, these phenotypes were not suppressed by *spt5*(*T1A*)*_7_*, which is expected to interfere with sense transcription-promoting but not –repressive effects of Cdk9 (22,26,61).

To account for a different kind of antagonism between Cdk9 and H2Bub1—their opposing effects on RNAPII distribution on transcribed genes—we previously proposed that H2Bub1 and Cdk9 cooperate to regulate RNAPII elongation in a rheostat-like mechanism: H2Bub1 provides both resistance to polymerase transit and a signal for the recruitment of Cdk9 to boost elongation rate while promoting further H2Bub1 formation through recruitment of PAF, Rhp6 (ortholog of Rad6) and Brl2 (28). We now propose that this enforced co-localization of H2Bub1 and Cdk9-dependent phosphorylations also serves to concentrate Clr6-CII activity where it is needed—at or near TECs—to deacetylate histone tails and suppress cryptic initiation within coding regions. In specific genomic contexts, however, Cdk9 and H2Bub1 pathways can transduce different regulatory inputs to exert opposing effects on gene expression (Fig. 7).

## Methods

### Yeast strains and growth conditions

Yeast strains used in this study are listed in Supplementary Table 3. Strains JTB413 (*alp13Δ::kanMX4*) and JTB414 (*cph1Δ::kanMX4*) were derived from *h^+^/h^+^* diploids (purchased from Bioneer) after transformation with plasmid pON177 to induce sporulation (62). Hygromycin-resistant versions of these strains (JTB621 and JTB622) were derived by PCR-based marker switch as described (63). The *htb1-K119R::hphMX6* allele was similarly derived from strain JTB86-3 (10). Correct integration of the markers was confirmed by diagnostic PCR. Double and triple mutant strains were constructed by tetrad dissection. Strains were cultured in YES (5% yeast extract, 30% dextrose, 250 mg/L each of adenine, leucine, histidine, and uracil) at 30^o^C. Septation phenotypes were scored on cells stained with DAPI and calcofluor as described previously (27).

### RNA-seq analysis

*S. pombe* cell cultures (50 mL) of strains JTB362 (WT), JTB425 (*cdk9^as^*), JTB97 (*htb1-K119R*), and JTB508 (*cdk9^as^ htb1-K119R*) were grown to OD_600_ of 0.2 before treatment with DMSO or 20 μM 3-MB-PP1 for 1 h. Total RNA was extracted by a hot phenol method (24) and further purified on Qiagen RNeasy columns. Strand-specific RNA-seq libraries were prepared from poly(A)-enriched RNA as described (64). Fastq files were aligned to the *S. pombe* ASM294v2 genome assembly. Resulting Sam-files (available on NCBI BioProject ID:382240) were uploaded to Podbat (65) where the data were further analyzed. For feature annotation, ASM294v2.20 was used. Data were normalized reads per kilobase per million (RPKM) and averaged over duplicates. Sense transcripts were defined as those that map to annotated protein-coding genes (5,123 loci), and antisense transcripts were defined as those that map antisense to annotated protein-coding genes (those that overlapped more than one gene were discarded). Intergenic transcripts were defined as those that map between annotated coding and non-coding loci. RPKM for each feature was calculated separately for sense and antisense. To determine differentially expressed genes, a cut-off of 2-fold (values in log 2 space >1 or <-1), with the highest absolute value being above 1 RPKM to filter out noise. Metagene analysis was performed in Podbat using normalized data. Comparison of RNA-seq data to previous strand-specific expression data was carried out as described (33). Clustering (hlcust) and chromosome position analysis was done in R.

### Quantitative RT-PCR

For strand-specific RT-qPCR to measure levels of *cdc2^+^* and *erg32^+^* antisense transcripts, 1-5 μg total RNA was converted to cDNA using the RevertAid H minus kit (Invitrogen) and the gene-specific primers listed in Supplementary Table 2. Expression levels were normalized to those obtained for the *act1^+^* sense transcript in each strain. To measure expression of subtelomeric genes and *tlh1^+^*, cDNA was synthesized from 1 μg RNA with an oligo-dT primer. Expression levels were normalized to *act1^+^* in each strain. Primers for qPCR are listed in Supplementary Table 4.

### Chromatin immunoprecipitation (ChIP)

ChIP was carried out as described previously (22,66). Briefly, *S. pombe* cell cultures were grown in YES to OD_600_ of 0.3-0.6, treated with DMSO or 20 μM 3-MB-PP1 for 1 h, and crosslinked with 1% formaldehyde for 15 min at room temperature. To terminate the crosslinking reaction 2.5 M glycine was added to a final concentration of 125 mM for 5 min. Cells were pelleted by centrifugation, washed twice with 10 mL cold TBS (10 mM Tris pH 7.5, 150 mM NaCl), frozen on dry ice and stored at −80 ^o^C. Cell pellets from 50 mL cultures were resuspended in 0.4 mL FA lysis buffer [50 mM HEPES pH 7.6, 150 mM NaCl, 1mM EDTA, 1% Triton X-100, 0.1% sodium deoxycholate, 1mM PMSF, and protease inhibitor cocktail (Roche)] and lysed in a mini-bead beater (Biospec products) in the presence of glass beads (50 micron; Sigma) at 4 ^o^C for 2 min. Lysates were centrifuged at 16,100 x g_av_ for 15 min at 4 ^o^C. Pellets containing chromatin were resuspended in 1mL FA Lysis buffer and transferred to 15 mL polycarbonate tubes. Lysates were then sonicated for 20 min at 4^o^C (30 sec on, 30 sec off, output setting high) using a waterbath sonicator (Diagenode Bioruptor). Sonicated samples were then transferred to a new 1.5-mL eppendorf tube and centrifuged at 16,100 x g_av_ for 5 minutes at 4^o^C. 100 μL of supernatant was kept as the “input” fraction, and the remainder (~900 μL) was subjected to immunoprecipitation with antibodies against histone H3 (Abcam 1791), K9,14-acetylated histone H3 (Millipore 06-599), myc epitope (9E10, Sigma), or control IgG. After incubation at 4^o^C with mixing for at least 4 h, immune complexes were recovered using 15 μL protein A/G agarose beads (Thermo Fisher) and incubated for an additional hour. Beads were washed sequentially with 0.5 mL FA Lysis + 0.1% SDS, FA Lysis + 0.1% SDS + 500 mM NaCl, LiCl buffer (10 mM Tris pH 7.5, 1 mM EDTA, 250 mM LiCl, 0.5% sodium deoxycholate, 0.5% NP-40), and TE (10 mM Tris pH 7.5, 1 mM EDTA). Each wash was for 4 min with mixing at room temperature. Immune complexes were eluted in 100 μL elution buffer (50 mM Tris pH 7.5, 10 mM EDTA, 1% SDS) at 65^o^C for 20 min. Beads were washed with 150 μL TE + 0.67% SDS, which was combined with the previous eluate. 150 μL TE + 0.67% SDS was also added to the input samples, and both IP and input samples were incubated at 65^o^C overnight to reverse protein-DNA crosslinks. DNA was purified by phenol/chloroform extraction as described previously (22). Analysis by qPCR was carried out using a Bio-Rad CFX96 instrument, Bio-Rad iQ Green SYBR mix, and the primers listed in Supplementary Table 2. ChIP signals were calculated as IP/input and normalized as indicated in the figure legends.

For ChIP-chip analysis, purified input and IP samples were amplified, labeled, and hybridized to *S. pombe* tiling microarrays containing 4x44K probes (Agilent Technologies) as described previously (22). Arrays were scanned on an Axon GenePix 4000B scanner and datasets are available on NCBIGEO (dataset GSE31071). Raw data were processed and converted to BED files as described previously. The histone H3 (K9,K14) acetylation level (normalized to histone H3) for each protein-coding gene was determined using Podbat software (65).

## Acknowledgments

We thank Chao Zhang (USC) and Kevan Shokat (UCSF) for providing 3-MB-PP1, Olaf Nielsen for providing plasmid pON177, and members of the Fisher and Tanny labs for helpful discussions. This work was supported by NIH grant GM104291 to R.P.F., Canadian Institutes of Health Research grants to J.C.T. (MOP-130362) and F.R. (MOP-133648), a Wellcome Trust Senior Investigator Award to J.B. (grant 095598/Z/11/Z), and Swedish Research Council (VR2015-02312) and Cancerfonden (CAN2016/576) awards to P.S. J.C.T. is supported by a fellowship from Fond de recherche Quebec Santé (33115).

